# Notch1 switches progenitor competence in inducing medulloblastoma

**DOI:** 10.1101/2020.05.10.084335

**Authors:** Claudio Ballabio, Matteo Gianesello, Chiara Lago, Konstantin Okonechnikov, Marica Anderle, Giuseppe Aiello, Francesco Antonica, Tingting Zhang, Francesca Gianno, Felice Giangaspero, Bassem A. Hassan, Stefan M. Pfister, Luca Tiberi

## Abstract

The identity of the cell of origin is a key determinant of cancer subtype, progression and prognosis. Group 3 Medulloblastoma (MB) is a malignant childhood brain cancer with poor prognosis and unknown cell of origin. We overexpressed the Group 3 MB genetic drivers MYC and Gfi1 in different candidate cells of origin in the postnatal mouse cerebellum. We found that S100b^+^ cells are competent to initiate Group 3 MB, while Math1^+^, Sox2^+^ or Ascl1^+^ cells are not. We noted that S100b^+^ cells have higher levels of Notch1 pathway activity compared to Math1^+^ cells. Interestingly, we found that additional activation of Notch1 in Math1^+^ cells was sufficient to induce Group 3 MB upon MYC/Gfi1 expression. Taken together, our data suggest that the MB cell of origin competence depends on the cellular identity, which relies on Notch1 activity.

**Graphical Abstract:** 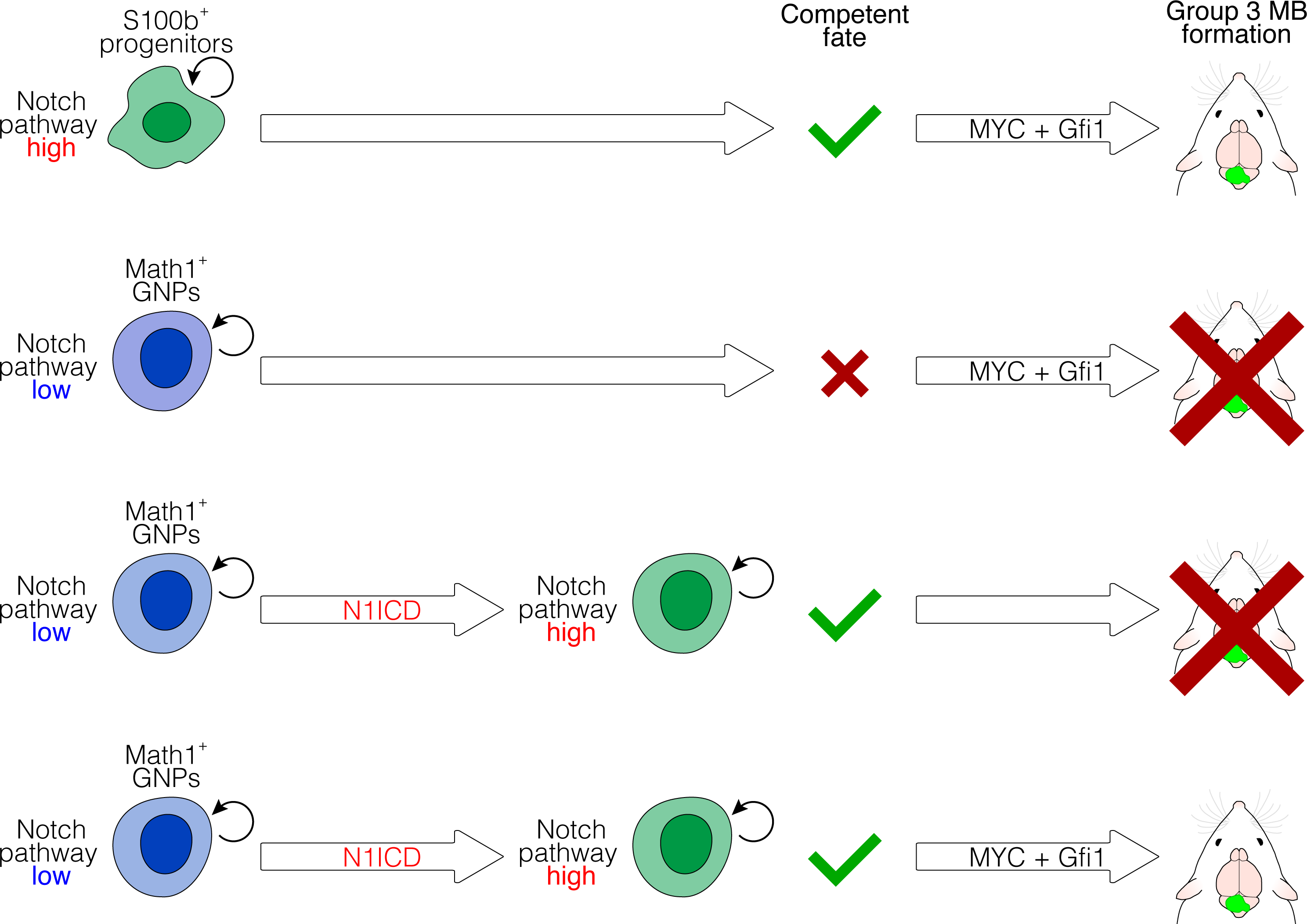

## Introduction

Defining the cancer cell of origin can be critical for understanding the first steps of cancer development and discovering the signals required for transformation (Gupta et al., 2019, Lytle et al., 2018). The identity of certain classes of tumors seems to be more strongly related to the cell of origin than to the oncogenic insult that induces malignant transformation. For example, in brain tumors concurrent inactivation of Trp53, Nf1 and Pten in neural progenitor cells or oligodendrocyte progenitors triggers the development of different subtypes of glioblastoma with distinct gene expression profiles (Alcantara et., 2009, 2015). BCR–ABL also provides an interesting example of an oncogene that produces different tumors depending on the cell in which it is expressed (Zhao et al., 2007). These studies suggest that the transcriptional context of the cell of origin can determine the identity of the tumor. By contrast, in some cases certain driver mutations rather than the cell of origin, mainly define the tumour profile. Indeed, activation of Hedgehog signalling in either neural stem cells, granule neural precursors or postmitotic granule neurons leads to development of aggressive medulloblastomas with similar molecular profiles (Aiello et al., 2019, Yang et al., 2008, Schuller et al., 2008). Nevertheless, it remains partially unclear what the specific determinants of the cell of origin required for tumor initiation are, and whether these specific features could be activated by an oncogenic insult only. For instance, the cell of origin of Group 3 medulloblastoma is currently unknown, but several genetic mutations able to generate this kind of tumors have been identified (Ballabio et al., 2020, Northcott et al., 2014). Group 3 is the most aggressive subtype of medulloblastoma (MB), mainly affecting children younger than 10 years of age. Recently, single cell RNA sequencing (scRNAseq) studies allowed the comparison of human cerebellar tumors with transcriptional clusters in the developing murine cerebellum. Analysis of human Group 3 MB revealed the presence of different transcriptional clusters within the tumor, resembling granule neuron precursors (GNPs), unipolar brush cells, Purkinje cells and GABAergic interneuron lineages during normal development (Hovestadt et al., 2019, Vladoiu et al, 2019). This suggested that early uncommitted human cerebellar stem cells might represent a putative Group 3 MB cell of origin (Vladoiu et al, 2019). The human cerebellum contains 80% of all brain neurons and has a 750-fold larger surface area, increased neuronal numbers, altered neuronal subtype ratios and increased folial complexity compared to the mouse cerebellum (Haldipur et al., 2019, Van Essen et al., 2002). Human cerebellar development begins 30 days after conception and is thought to be completed by the age of 2 years (Haldipur et al., 2019). Human cerebellar volume increases five-fold between 22 post-conception weeks (PCW) and birth and it becomes highly foliated during the third trimester (24 to 40 gestational weeks) (Haldipur et al., 2019). Granule cell progenitor proliferation peaks during this period and it is accompanied by increased external granule layer thickness. In mice, cerebellar growth and foliation are driven by granule cell progenitor proliferation between postnatal day 1 (P1) and P14, with deficient proliferation causing external granule layer thinning (Butts et al., 2014). Hence, the postnatal development of the mouse cerebellum resembles key aspects of human embryonic development. For this reason, the cell of origin of Group 3 MB should also be studied during postnatal mouse cerebellar development. Indeed, MB mouse models have been developed by postnatally deregulating *Myc* and *Trp53 ex vivo* in Math1^+^ GNPs or in CD133^+^ stem cells (Kawauchi et al., 2012) (Pei et al., 2012). It has also been demonstrated that CD133^+^ cerebellar stem cells are able to give rise to Group 3 MB when transduced *ex vivo* with *Myc* and *Gfi1* or with *My c* a n d *Gfi1b* (Northcott et al., 2014, Lee et al., 2019). Furthermore, we have recently demonstrated that MYC and Gfi1 induce Group 3 MB when co-overexpressed in human brain organoids (Ballabio et al., 2020). Moreover, Sox2^+^ astrocyte progenitors (upon Myc overexpression *ex vivo*) have been proposed to give rise to Group 3 MB in the postnatal developing cerebellum (Tao et al., 2019). Notably, whether MYC and Gfi1 induce Group 3 MB in specific progenitor populations has never been tested *in vivo*, during embryonic or postnatal cerebellum development.

## Results

### S100b-positive cells are competent to induce Group 3 MB

In order to modify mouse cerebellar cells directly *in vivo*, we stably transfected postnatal day 0 (P0) mouse cerebella exploiting the PiggyBac transposase system. We were able to target different cell populations in the developing cerebellum, such as Pax6^+^, Sox2^+^, Sox9^+^ and Calbindin^+^ cells (Figure S1A,B)(Ballabio et al., 2020). To investigate which postnatal progenitors/cells are competent to potentially give rise to Group 3 MB, we used mice expressing creER recombinase in GNPs (Math1-creER, 24 mice)(Machold et al., 2005), in glial cells progenitors (Sox2−creER, 7 mice)(Arnold et al., 2011), Purkinje cells, GABAergic interneurons and glial cells (Ascl1-creER, 24 mice)(Sudarov et al., 2011)(Figure 1A, Figure S1C). We crossed these mice with R26-LSL-Myc transgenic mice conditionally expressing MYC under the control of the creER recombinase and we stably transfected the cerebella at P0 with PiggyBac vectors expressing Gfi1 (pPB-Gfi1). In parallel, we transfected the cerebella of mice bearing creER recombinase with PiggyBac vectors expressing both MYC and Gfi1 under the control of a loxP-STOP-loxP cassette (pPB-LSL-MYC, pPB-LSL-Gfi1) at P0 (Figure 1A). Subsequently, we induced the creER-dependent removal of the LSL cassette by injecting tamoxifen (TMX) at different time points (Table S1, we considered only mice with transfected cells). To further identify which progenitors are competent to give rise to Group 3 MB, we also transfected pPB-LSL-MYC and pPB-LSL-Gfi1 vectors together with plasmids expressing the cre recombinase under the control of *Tbr2* (UBC cells), *Sox2*, *Math1* or *S100b* (astrocytic and extra-astrocytic expression in human brain) (Steiner et al., 2007) regulatory elements (Figure 1B, Table S1). Notably, we obtained tumors only when MYC and Gfi1 expression was driven by *S100b* promoter (3 out of 16, from two different litters, Figure 1C). These tumors showed few S100b^+^ cells (Figure 1D), consistently with the observation that the tumor phenotype not always resembles its cell of origin (Proia et al., 2011, Van Keymeulen et al., 2015). In addition, these tumors were pH3^+^, showed few GFAP^+^ cells, and were also positive for the Group 3-specific marker NPR3 (Figure 1E, Figure S1D,E). Interestingly, Sox2−creER;R26-LSL-Myc mice developed choroid plexus carcinoma (CPC) 1 month after tamoxifen injection (6 out of 6, Figure 1F and data not shown) as already published (Kawauchi et al., 2017). We did not detect MB formation with any of the other promoters, but we observed Venus^+^ cells 75 days after injection (Figure 1G,H, Figure S1F,G). We checked the expression of MYC and Gfi1 by immunofluorescence (Figure S1H-J). Indeed, we were able to detect the expression of MYC in the CPC developed by Sox2−creER;R26-LSL-Myc transgenic mice injected with tamoxifen at P2 (Figure S1H). The expression of both Gfi1 and MYC was also observed by immunofluorescence in P7 CD1 mouse cerebella transfected at P0 with pMath1-cre + pPB-LSL-Myc + pPB-LSL-Gfi1 (Figure S1I,J). Since the *S100b* promoter was able to drive cre expression in cells competent to develop Group 3 MB, we investigated the identity of these cells. As shown in Figure S2A, S100b^+^ cells are present in the mouse cerebellum at P0. In addition, we could detect Venus^+^/S100b^+^ cells 4 days after transfection with pS100b-cre + pPB-LSL-Venus, thus confirming that our promoter recapitulates endogenous S100b expression (Figure 2A,B). Furthermore, *S100b* promoter drives Venus expression mostly in the ventricular zone (VZ), but also in the white matter (WM), internal granule layer (IGL), molecular layer (ML) and external granule layer (EGL) and most of those cells are positive for Sox2 and GFAP (Figure 2C,D, glial cells). Since the Sox2 promoter is not able to render postnatal cells competent for Group 3 MB development, we investigated the presence of S100b^+^ (Venus^+^) and Sox2^−^cells (Figure 2D,E, S2B,C). Indeed, we observed that 34.8% ± 7.7% (mean ± s.e.m., *n* = 7 brains) of S100b-cre^+^ cells are Sox2^−^4 days after transfection, with a small subset being Sox2^−^/Nestin^+^ (Figure 2D). We then characterized which cells are produced by S100b-cre^+^ cells by performing lineage tracing experiments. 30 days after transfection, we observed ependymal cells (Figure 2F), glial cells (Figure S2D), oligodendrocytes (Figure 2G), Bergmann glia (Figure 2H), and S100b^+^ cells (Figure 2I) that derive from S100b-cre^+^ cells transfected at P0. Hence, the population of S100b-cre ^+^ cells at P0 are able to generate several cerebellar populations, notably different types of glial cells. To study the first step of S100b cell transformation we performed lineage tracing of S100b-cre ^+^ cells that start to stably express MYC and Gfi1 starting from P0. As shown in Figure 2J-M, after 10 days we observed small clusters of Venus^+^ cells, having a homogeneous round morphology that is different from cells not expressing MYC and Gfi1 (Figure 2E, S2B,C). Furthermore, these cell clusters are mainly GFAP and S100b negative (Figure 2J,M), with few Sox2^+^ and Sox9^+^ cells (Figure 2K,L). Remarkably, we did not detect any cluster formation with other promoters driving cre expression (Figure 1G,H, S1F,G, data not shown). Since S100b^+^ cells are able to generate Group 3 MB postnatally, we tested whether S100b^+^ cells might be competent to give rise to MB also during embryonic development. We first confirmed the embryonic expression of S100b in the cerebellar ventricular zone (Figure S2E) (Hachem et al., 2007) and then we electroporated *in utero* pPB-LSL-Myc + pPB-LSL-Gfi1 together with pS100b-cre at embryonic day E15.5 (Figure S2F). Interestingly, we observed formation of tumors in 3 out of 12 electroporated mice (Figure S2G) that were positive for the Group 3-specific marker NPR3 (Figure S2H).

**Figure 1.**
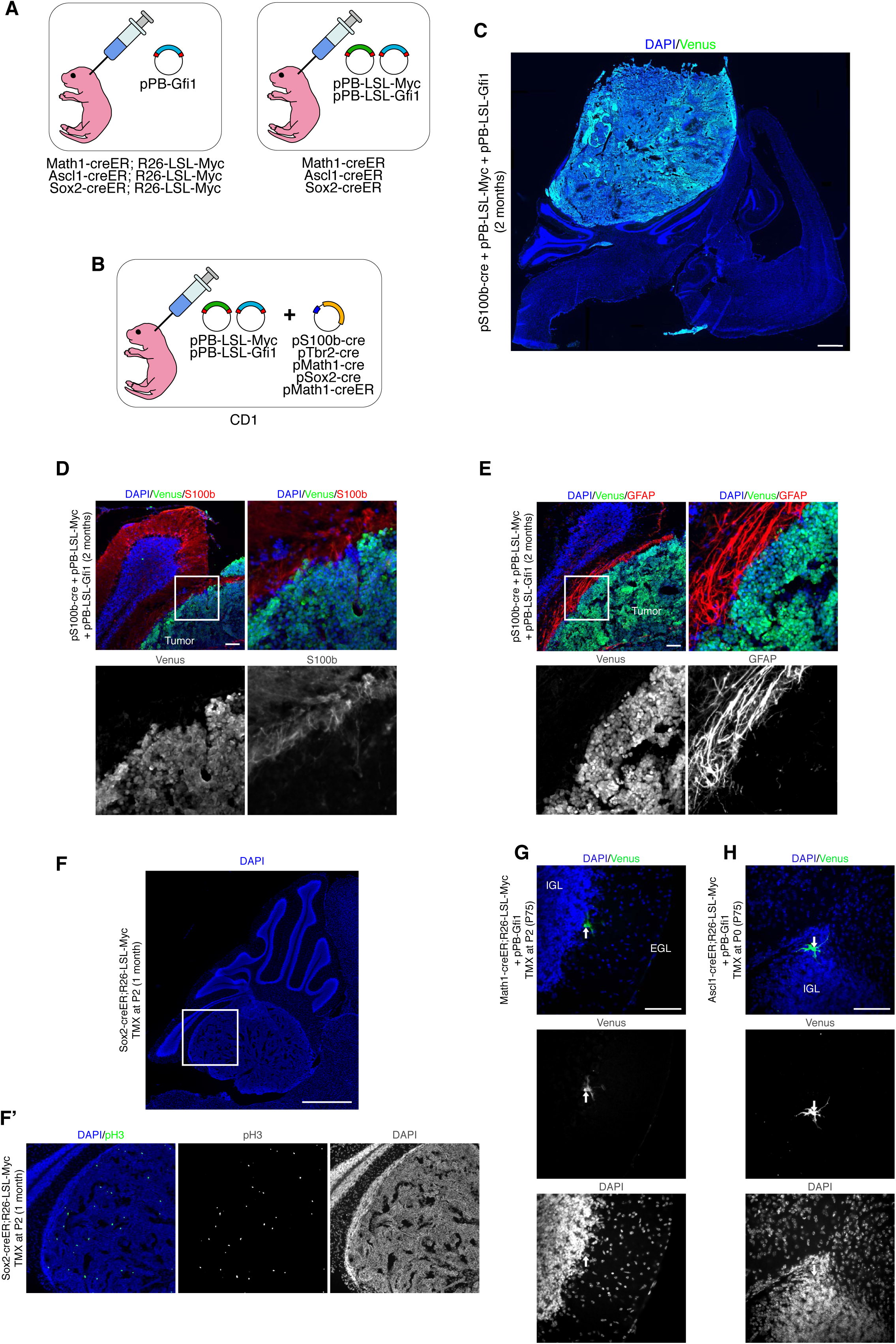
Overexpression of MYC and Gfi1 in postnatal S100b^+^ cerebellar cells induces MB. (**A,B**) Schematic representation of *in vivo* transfection experiments to overexpress Myc and Gfi1 in different postnatal cerebellar cell populations. (**C**) DAPI staining and Venus immunofluorescence of sagittal brain section of CD1 mouse 2 months after transfection with pPBase + pS100b-cre + pPB-LSL-Myc + pPB-LSL-Gfi1 + pPB-LSL-Venus at P0. (**D,E**) Confocal images of Venus and S100b (**D**), Venus and GFAP (**E**) immunofluorescence of tumors in CD1 mice 2 months after transfection with pPBase + pS100b-cre + pPB-LSL-Myc + pPB-LSL-Gfi1 + pPB-LSL-Venus at P0. The white squares in (**D,E**) mark the regions shown at higher magnification. (**F**) DAPI staining and pH3 immunofluorescence of sagittal brain section of Sox2-creER;R26-LSL-Myc mouse 1 month after tamoxifen injection at P2. The white square in (**F**) marks the region shown in (**F’**). (**G,H**) DAPI staining and Venus immunofluorescence of sagittal brain sections of Math1-creER;R26-LSL-Myc mouse (**G**) or Ascl1-creER;R26-LSL-Myc mouse (**H**) 2.5 months after transfection with pPBase + pPB-Gfi1 + pPB-Venus at P0 and tamoxifen injection at P2 (**G**) or P0 (**H**). Arrows point to Venus^+^ cells. Scale bars: 1 mm in (**C,F**), 100 µm in (**D,E,G,H**).

**Figure 2.**
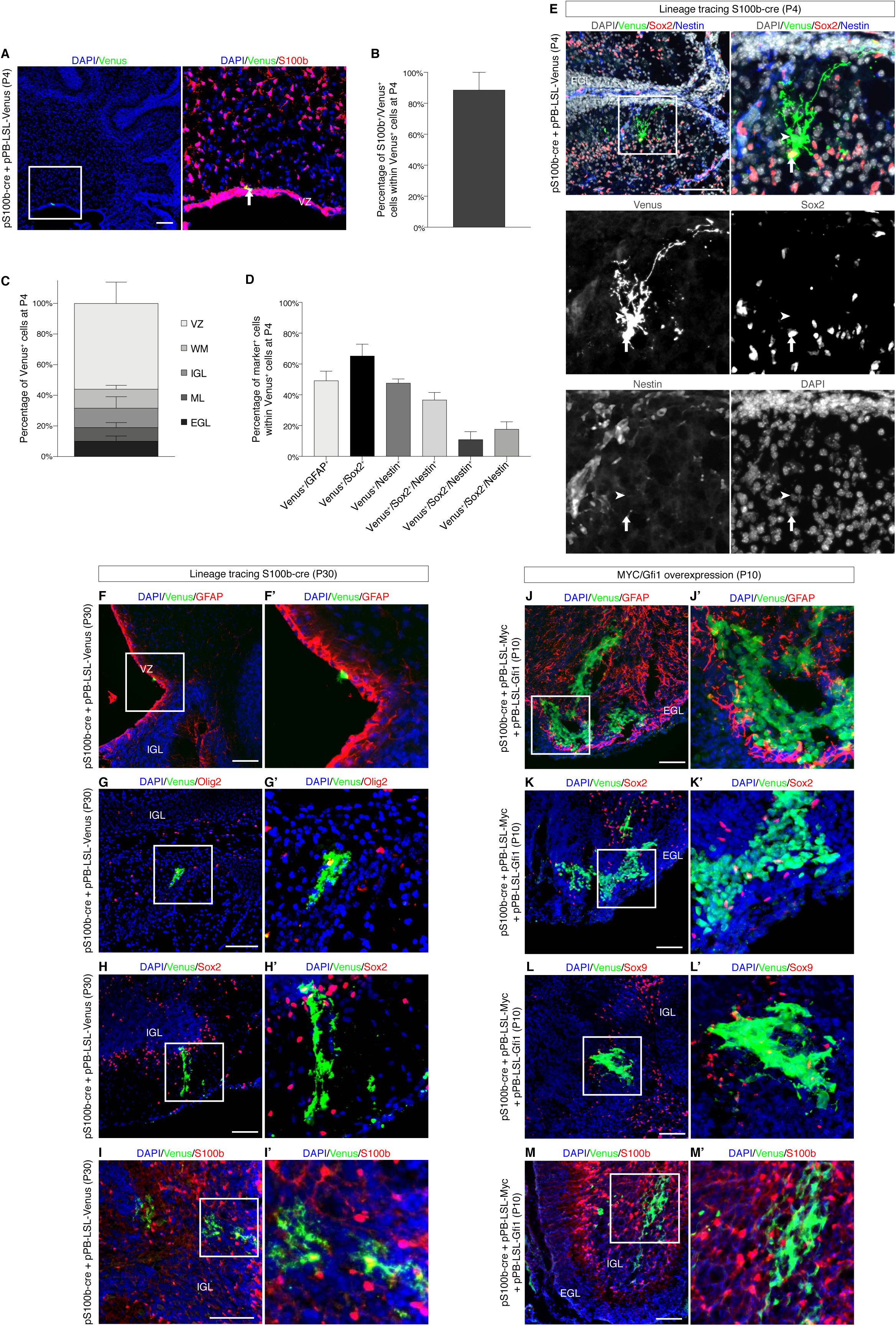
Lineage tracing of S100b-cre^+^ cells in postnatal mouse cerebellum. (**A**) Confocal images of Venus and S100b immunofluorescence of CD1 mouse cerebellum 4 days after transfection with pPBase + pS100b-cre + pPB-LSL-Venus at P0. The white square marks the region shown at higher magnification. The arrow points to a Venus^+^/S100b^+^ cell. (**B**) Quantification of Venus^+^/S100b^+^ cells within Venus^+^ cells in CD1 mouse cerebellum 4 days after transfection with pPBase + pS100b-cre + pPB-LSL-Venus at P0 (n=2 mice, 99 cells). (**C**) Quantification of Venus^+^ cells within different regions of CD1 mouse cerebellum 4 days after transfection with pPBase + pS100b-cre + pPB-LSL-Venus at P0 (n=5 mice, 362 cells). VZ: ventricular zone; WM: white matter; IGL: internal granule layer; ML: molecular layer; EGL: external granule layer. (**D**) Quantification of Venus^+^/GFAP^+^ cells (n=3 mice, 99 cells), Venus^+^/Sox2^+^ cells (n=7 mice, 188 cells), Venus^+^/Nestin^+^ cells (n=5 mice, 81 cells), Venus^+^/Sox2^+^/Nestin^+^ cells (n=5 mice, 81 cells), Venus^+^/Sox2^−^/Nestin^+^ cells (n=5 mice, 81 cells) and Venus^+^/Sox2^−^/Nestin^-^cells (n=5 mice, 81 cells) within Venus^+^ cells in CD1 mouse cerebellum 4 days after transfection with pPBase + pS100b-cre + pPB-LSL-Venus at P0. (**E**) Confocal images of Venus, Sox2 and Nestin immunofluorescence of CD1 mouse cerebellum 4 days after transfection with pPBase + pS100b-cre + pPB-LSL-Venus at P0. The white square marks the region shown at higher magnification. The arrow points to a Venus^+^/Sox2^+^ cell, whereas the arrowhead points to a Venus^+^/Sox2^−^ cell. (**F-I**) Confocal images of Venus and GFAP (**F**), Venus and Olig2 (**G**), Venus and Sox2 (**H**), Venus and S100b (**I**) immunofluorescence of CD1 mouse cerebellum 30 days after transfection with pPBase + pS100b-cre + pPB-LSL-Venus at P0. The white squares in (**F-I**) mark the regions shown at higher magnification in (**F’-I’**) respectively. (**J-M**) Confocal images of Venus and GFAP (**J**), Venus and Sox2 (**K**), Venus and Sox9 (**L**), Venus and S100b (**M**) immunofluorescence of CD1 mouse cerebellum 10 days after transfection with pPBase + pS100b-cre + pPB-LSL-Myc + pPB-LSL-Gfi1 + pPB-LSL-Venus at P0. The white squares in (**J-M**) mark the regions shown at higher magnification in (**J’-M’**) respectively. Scale bars: 100 µm in (**A,E-M**). Data in (**B-D**) are presented as mean + s.e.m.

### Human S100B-positive cells are competent to induce Group 3 MB

To test whether S100b cells are also present during human cerebellar development we performed histological analysis of S100B in human tissues. As shown in Figure 3A,B, S100B^+^ cells are present in the human cerebellum at 22 GW and 39 GW in EGL, IGL and ML. Interestingly, we could also detect a few S100B^+^ cells in human Group 3 MB samples (Figure S3A). Based on these results, we tested whether human S100B^+^ cells are competent to generate MB in human cerebellar organoids. Indeed, we have recently shown that MYC and Gfi1 overexpression in human cerebellar organoids induces Group3 MB with a methylation profile similar to human patients (Ballabio et al., 2020). Therefore, we electroporated human iPSC-derived cerebellar organoids (Ballabio et al., 2020, Muguruma et al., 2015) with pPB-LSL-Venus and pS100b-Cre or pSox2−Cre at day 35, in order to target S100B^+^ and SOX2^+^ cells when these cerebellar progenitors are present (Figure S3B-D) (Ballabio et al., 2020, Muguruma et al., 2015). Unfortunately, we were unable to specifically target GNPs in human cerebellar organoids, as reliable human ATOH1 regulatory sequences are lacking. Next, we also electroporated pPB-LSL-Myc + pPB-LSL-Gfi1 together with pS100b-Cre or pSox2−Cre. As shown in Figure 3C,D, MYC and Gfi1 expression in S100B-cre^+^ cells induced formation of clusters of cells in 12,5% of electroporated organoids (n=40) similarly as we previously observed (Ballabio et al., 2020). Interestingly, MYC and Gfi1 overexpression in Sox2−cre^+^ cells did not induce cell cluster formation (0 out of 40) (Figure 3C,D). Furthermore, we observed clusters of PCNA^+^ cells using the *S100B* promoter, suggesting that MYC and Gfi1 overexpression have an oncogenic potential in S100B-cre^+^ cells also in human cerebellar organoids (Figure 3E).

**Figure 3.**
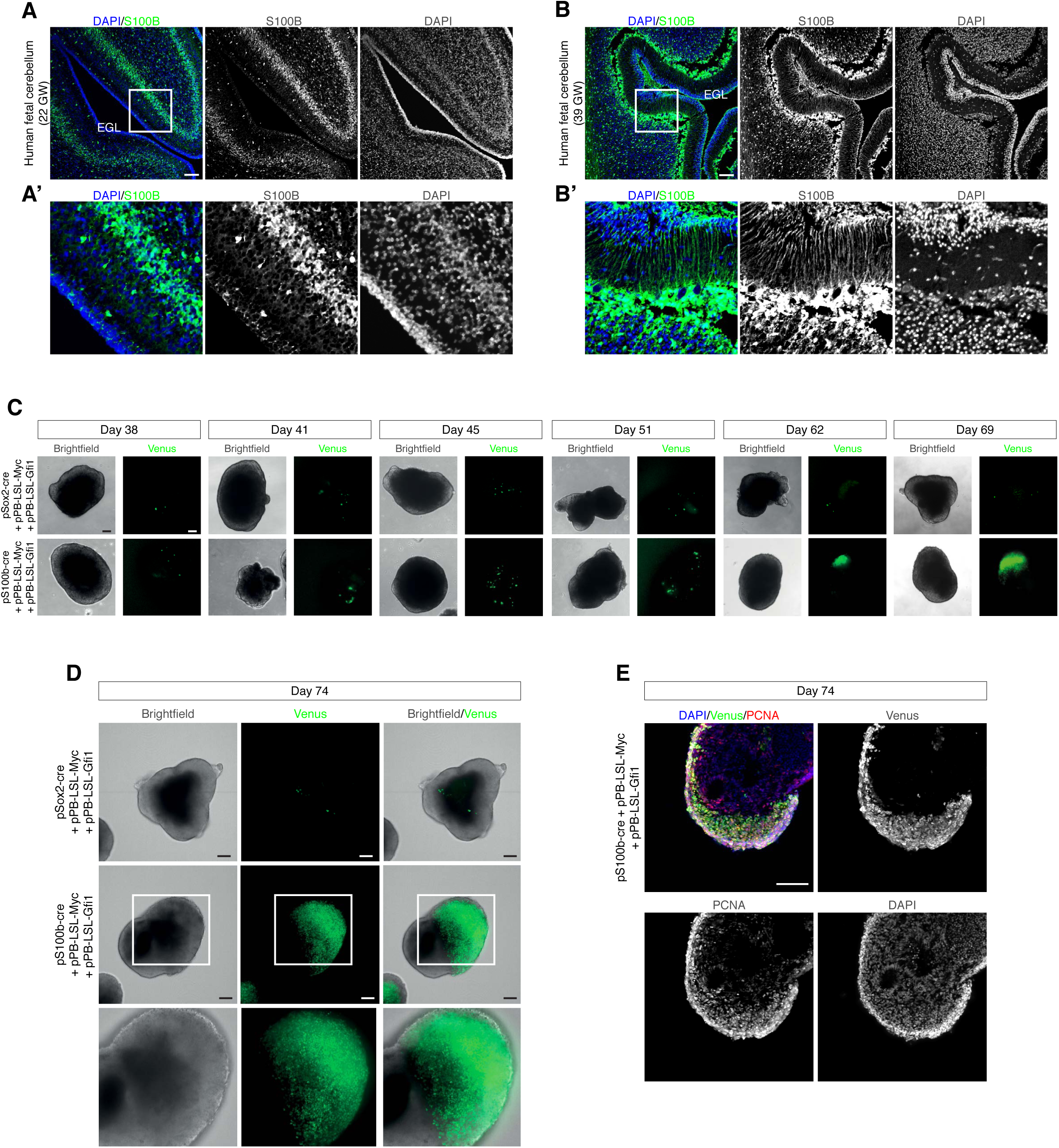
Human S100B-cre^+^ cells are responsive to MYC and Gfi1 overexpression. (**A**) Confocal images of S100B immunofluorescence of human fetal cerebellum at 22 GW (gestational week). (**B**) Confocal images of S100B immunofluorescence of human fetal cerebellum at 39 GW. The white squares in (**A,B**) mark the regions shown at higher magnification in (**A’,B’**) respectively. (**C**) Brightfield and fluorescence images of cerebellar organoids at different time points electroporated at day 35 with pPBase + pSox2-cre + pPB-LSL-Gfi1 + pPB-LSL-Myc + pPB-LSL-Venus, or pPBase + pS100b-cre + pPB-LSL-Gfi1 + pPB-LSL-Myc + pPB-LSL-Venus. (**D**) Brightfield and fluorescence images of cerebellar organoids at 74 days, electroporated at day 35 with pPBase + pSox2-cre + pPB-LSL-Gfi1 + pPB-LSL-Myc + pPB-LSL-Venus or pPBase + pS100b-cre + pPB-LSL-Gfi1 + pPB-LSL-Myc + pPB-LSL-Venus. The white squares mark the regions shown at higher magnification. (**E**) Confocal images of Venus and PCNA immunofluorescence of cerebellar organoids at day 74, electroporated at day 35 with pPBase + pS100b-cre + pPB-LSL-Gfi1 + pPB-LSL-Myc + pPB-LSL-Venus. Scale bars: 100 µm in (**A-E**).

### Notch1 activation makes Math1-positive progenitors competent to induce Group3 MB

Based on our data we speculated that S100b^+^ cells should possess specific features/pathways that are not present in postnatal Math1^+^, Ascl1^+^ and Sox2^+^ cells. Interestingly, it has been found that Notch signaling regulates fate decisions of early mouse cerebellar progenitors (Machold et al., 2007, Zhang et al., 2020). In particular, Sox2^+^ progenitors at E9.5 have high Notch1 pathway levels and are competent to give rise to Ascl1^+^ and Math1^+^ progenitors (Zhang et al., 2020). We tried to target these Sox2^+^ early progenitors conditionally by overexpressing Myc at E9.5 and at E13.5 (Sox2−creER;R26-LSL-Myc mouse), but we were not able to obtain live pups, possibly due to a widespread Sox2 expression during mouse development (n=2 pregnant females administered with Tamoxifen at E9.5; n=2 pregnant females administered with Tamoxifen at E13.5). Notably, these early Sox2^+^ progenitors do not generate the postnatal Sox2^+^ cells that are not competent to induce Group 3 MB (Sox2−creER;R26-LSL-tdTomato, Tamoxifen at E10.5, Figure S4A). Furthermore, embryonic Math1^+^ progenitors express low levels of Notch1 and Hes1/Hes5 target genes and Notch1 pathway activation is able to repress Math1 expression, thus changing the fate of these progenitors (Zhang et al., 2020). The Notch signaling pathway plays a critical role in CNS development, stem cell maintenance and differentiation of cerebellar granule neuron precursors; modulates epithelial-to-mesenchymal transition; anyhow, its role in SHH MB is still controversial (Fan et al., 2004, Hatton et al., 2010, Julian et al., 2010, Natarajan et al., 2013). Mutations in NOTCH signaling genes have been described in Group 3 MB (Northcott et al., 2017), with especially elevated expression of NOTCH1 in spinal metastases (Kahn et al., 2018).

Based on these data we tested whether the Notch1 pathway plays a role in determining the competence of the different cerebellar progenitors to generate Group 3 MB. Using available scRNAseq data (Carter et al, 2018) from postnatal (P0-P4) mouse cerebella, we verified the stable cell types assignment for the different cerebellar cell populations (Figure 4A, Figure S4B). Further analyses on the scRNAseq data indicated that S100b is expressed in clusters assigned to the astrocytic lineage (which includes glial progenitors, oligodendrocytes, glia and Bergmann glia; cluster β and γ) and ciliated cells (cluster α (Carter et al, 2018)(Figure 4A,B-D), and is mostly mutually exclusive with the expression of Math1 (Figure 4E-G), especially in the granule neurons lineage. Therefore, we asked whether the expression of Notch pathway genes correlates with any cell lineage also in the postnatal cerebellar scRNAseq data, as in the embryonic mouse cerebellum (Zhang et al., 2020). As shown in Figure 4H-P and Figure S4C,D, we found that the S100b^+^ cell clusters show the highest expression levels of the Notch1 receptor and of Hes1/Hes5 target genes. On the other hand, postnatal Math1^+^ cells display lower levels of Notch pathway activation, similar to the Math1^+^ progenitors in the embryonic mouse cerebellum (Zhang et al, 2020). To confirm these data, we tested Notch activity using a well-known reporter for Notch signaling. We used a plasmid that drives expression of destabilized enhanced green fluorescent protein (d2EGFP) under the control of the Hes5 promoter (Ohtsuka et al., 2006). The promoter activity is dependent on Notch pathway activation and is restricted to neural stem cells in the mouse brain (Ohtsuka et al., 2006, Tiberi et al., 2012). As shown in Figure S4E-G, we observed more d2EGFP expression in S100b^+^ cells compared to Math1^+^ progenitors confirming that the Notch pathway is more active in S100b^+^ cells. Since constitutive activation of Notch1 suppresses the generation of Math1^+^ progenitors (Zhang et al., 2020), we tested if increasing Notch pathway activity could make Math1^+^ cells competent to induce Group 3 MB. To do so, we overexpressed N1ICD in Math1^+^ cells, which alone did not induce MB formation (Figure 5A). Next, we co-overexpressed N1ICD with MYC and Gfi1 to induce Group 3 MB in Math1^+^ progenitors. As shown in Figure 5B, N1ICD+MYC/Gfi1 overexpression in Math1^+^ progenitors induces tumor formation in 13 out of 14 mice (Figure 5A). These tumors are GFAP-negative, pH3-positive (Figure S5A,B), and are comparable to human Group 3 MB and to previously published Group 3 MB mouse models (MYC/Gfi1 and MYC/Otx2, Figure 5C)(Ballabio et al., 2020). In order to study the effects of N1ICD on Math1^+^ progenitors, we checked the transfected cerebella 14 days after injection: we found that N1ICD^+^MYC/Gfi1 transfected cells already formed small clusters of Ki67^+^ cells in the EGL, unlike granule neuron progenitors transfected with N1ICD alone. (Figure 5D-G).

**Figure 4.**
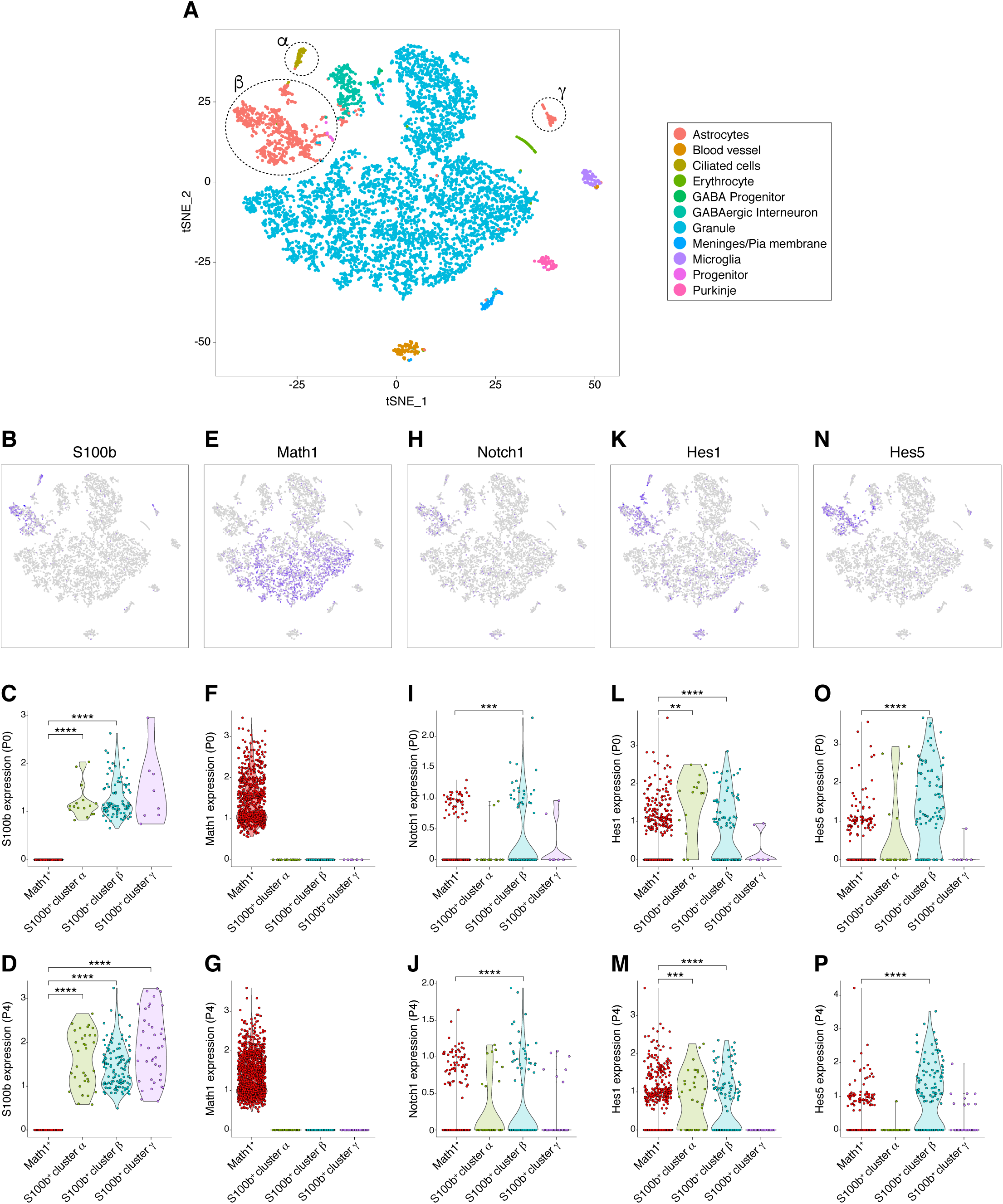
Notch pathway expression in the mouse postnatal cerebellum. (**A**) t-distributed stochastic neighbor embedding (t-SNE) visualization of cerebellar derived cell clusters at P0 and P4. Each point represents one cell. Main clusters are marked in color, while relevant subclusters are labelled as α, β and γ. (**B-P**) scRNAseq analysis of cell type specific markers in different cell clusters. (**B,E,H,K,N**) t-SNE visualization showing expression of cell type specific markers: S100b (**B**), Math1 (**E**), Notch1 (**H**), Hes1 (**K**), Hes5 (**N**). Cells are color-coded according to genes expression. (**C,F,I,L,O**) Violin plots to visualize gene expression levels of S100b (**C**), Math1 (**F**), Notch1 (**I**), Hes1 (**L**), Hes5 (**O**) at P0 in four groups: mutually exclusive Math1^+^ cells and S100b^+^ cells within cluster α, S100b^+^ cells within cluster β and S100b^+^ cells within cluster γ. (**D,G,J,M,P**) Violin plots to visualize gene expression levels of S100b (**D**), Math1 (**G**), Notch1 (**J**), Hes1 (**M**), Hes5 (**P**) at P4 in four groups: mutually exclusive Math1^+^ cells and S100b + cells within cluster α, S100b^+^ cells within cluster β and S100b^+^ cells within cluster γ. Unpaired Student’s *t* test. **q-value < 0.01. ***q-value < 0.001. ****q-value < 0.0001.

**Figure 5.**
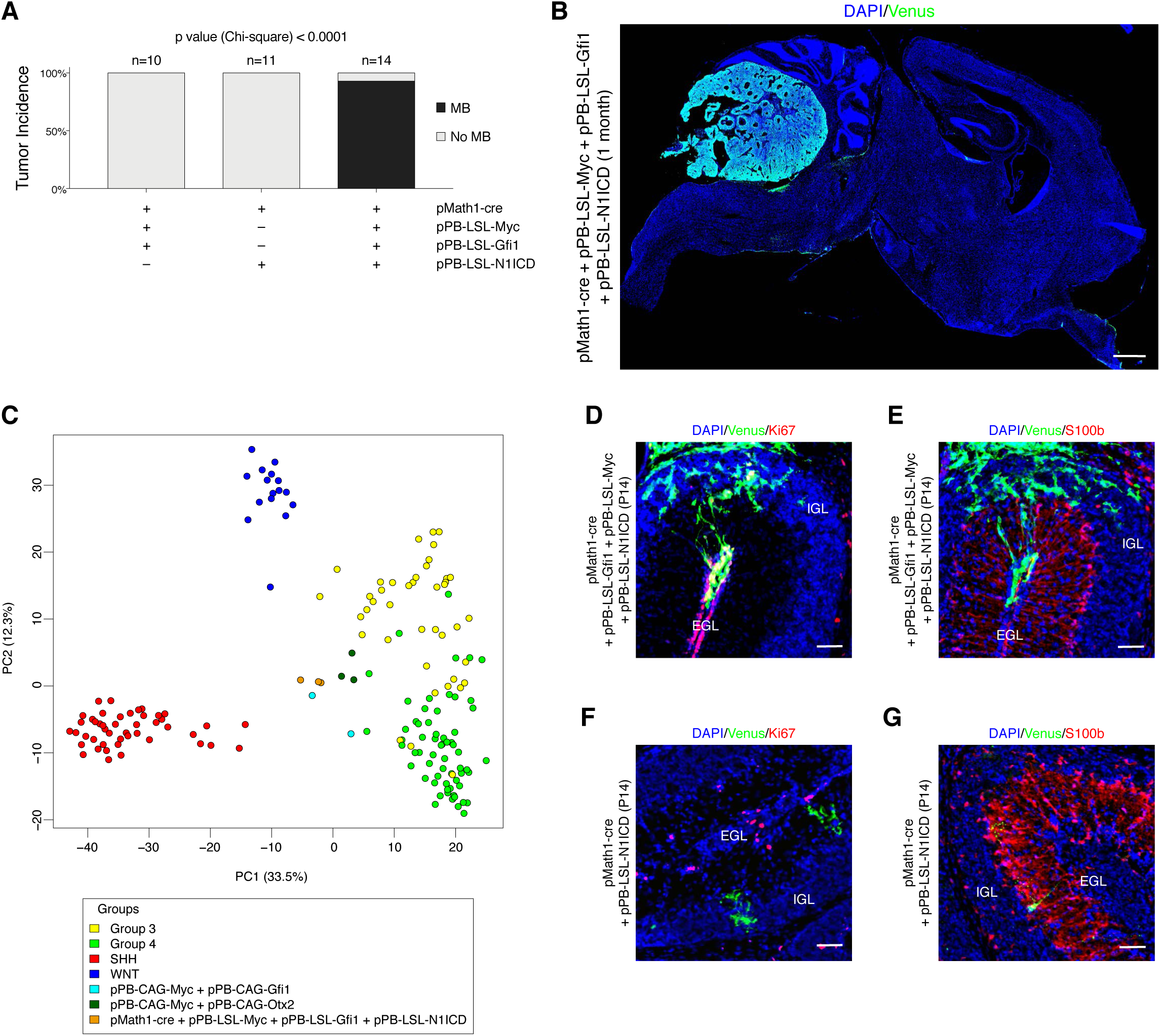
Notch pathway determines competence to generate Group 3 MB. (**A**) Histograms show the percentage of mice that develop MB (75 days) after transfection at P0 with either pPBase + pMath1-cre + pPB-LSL-Myc + pPB-LSL-Gfi1 + pPB-LSL-Venus, pPBase + pPB-LSL-N1ICD + pPB-LSL-Venus, or pPBase + pMath1-cre + pPB-LSL-Myc + pPB-LSL-Gfi1 + pPB-LSL-N1ICD + pPB-LSL-Venus. (**B**) DAPI staining and GFP (Venus) immunofluorescence of CD1 mouse brain sagittal section 1 month after transfection with pPBase ^+^ pMath1-cre + pPB-LSL-Myc + pPB-LSL-Gfi1 + pPB-LSL-N1ICD + pPB-LSL-Venus at P0. (**C**) Principal component analysis (PCA) visualization based on the combination of transcriptional profiles with orthologous genes control for the four consensus human MB subgroups together with previously published Group 3 MB mouse models (pPB-CAG-Myc + pPB-CAG-Gfi1; pPB-CAG-Myc + pPB-CAG-Otx2) and tumors developed after transfection of pPBase + pMath1-cre + pPB-LSL-Myc + pPB-LSL-Gfi1 + pPB-LSL-N1ICD + pPB-LSL-Venus. (**D-G**) Confocal images of GFP (Venus) and Ki67 (**D,F**) or S100b (**E,G**) immunofluorescence of transfected cells in CD1 mice 14 days after transfection with pPBase + pMath1-cre + pPB-LSL-Myc + pPB-LSL-Gfi1 + pPB-LSL-N1ICD + pPB-LSL-Venus (**D,E**), or pPBase + pMath1-cre + pPB-LSL-N1ICD + pPB-LSL-Venus (**F,G**) at P0. Chi-square test (**A**). Scale bars: 1 mm in (**B**), 50 µm in (**D-G**).

## Discussion

Despite large effort in understanding the cell of origin of different tumors, the relationship of the contribution of the cell of origin versus the relevant driver mutations is still elusive. Current opinion in the field is that tumor identity can be defined by the cell of origin and/or genetic mutation (Gupta et a., 2019, Lytle et al., 2018). In the first scenario, the cells of origin possess specific features, such a s the epigenetic state, that make them competent in inducing cancer and may also direct cell tumour-initiating capacity. Recent studies have shown that the epigenetic landscape is a critical determinant of both transformation susceptibility and the acquisition of a stem or progenitor fate (Xie et al., 2018, Yu et al., 2018). For example, work in zebrafish has shown that the epigenetic state can act as a determining factor for the cell of origin of melanoma (Kaufman et al., 2016). On the other hand, in medulloblastoma tumors, SHH activation produces the same tumors starting from different cells (Aiello et al., 2019, Yang et al., 2008, Schuller et al., 2008). Furthermore, in breast cancer oncogenic insults can change cell fate, and this fate is reflected in the tumor phenotype (Proia et al., 2011, Van Keymeulen et al., 2015). Notably, these oncogenic cues have been modulated in cells that are competent to induce cancer and change their cellular fate before the tumor initiation. It is possible that certain mutations are powerful enough in terms of defining cell fate, being able to override the transcriptional context of the cell of origin. Our work shows that S100b^+^ cells are competent to induce medulloblastoma while Math1^+^, Sox2^+^ or Ascl1^+^ cells seem to be unresponsive to MYC/Gfi1 overexpression. Interestingly, one of the molecular differences between these cells is that S100b^+^ cells present high levels of Notch pathway activity. By overexpressing the active Notch1 form (N1ICD), we demonstrated that Notch1 pathway activation, upon MYC/Gfi1 overexpression, induces cancer in Math1^+^ cell, that normally are not competent to induce Group 3 medulloblastoma. Therefore, we speculate that the levels of Notch1 signaling are critical for determining the fate of the Math1^+^ cells as previously described by Zhang et al., 2020. Moreover, we demonstrated that this fate switch is sufficient to allow the tumorigenesis. Notably, Notch1 overexpression *per se* is not sufficient to induce cancer in Math1^+^ progenitors but requires the overexpression of MYC/Gfi1, as in S100b^+^ cells. Based on our data, we speculate that previously described cell populations not able to originate human tumors should be challenged with activation of cell fate pathways. Notably, in human cerebellum S100B is expressed in many neural cell-types, progenitors and is less “astrocyte-specific” than GFAP (Steiner et al., 2007), suggesting that considering S100B as an astrocyte-specific marker might not be correct. We validated our findings also in human cerebellar organoids and our experiments clarified that S100B-cre ^+^ cells are competent to induce hyper-proliferation in cerebellar organoids, while Sox2−cre^+^ cells are not. Interestingly, MB mouse models have been already developed by postnatally deregulating *Myc* and/or Gfi1 *ex vivo* in Math1^+^ and Sox2^+^ cells (Tao et al, 2019, Kawauchi et al, 2012). Notably, cell dissociation for FACS sorting and *ex vivo* manipulation is able to activate Notch pathway (Rand et al., 2000, Liu et al., 2014) and, in light of our data, it appears to be a determinant of cell competence. Taken together, our data point to the direction that a subpopulation of S100b^+^ cells with high levels of Notch1 pathway might represent a good candidate as a cell of origin of Group 3 MB. More broadly, the involvement of Notch activity in determining progenitor identity in a wide variety of contexts suggests that this pathway may be a general regulator of cancer cell of origin competence.

## Author Contributions

C.B., M.G., G.A., performed *in vivo* experiments. C.L., M.A., performed organoids experiments and live imaging. F.A. performed live imaging. C.B., M.G., and L.T. designed and analyzed all experiments and wrote the manuscript. Fe.G., Fr.G., provided the human cerebellar tissue. T.Z performed the lineage tracing experiments with Sox2-creER mice at E10.5 supervised by B.H. K.O. analyzed the mouse cerebellar scRNA-seq data and the tumor RNA-seq data supervised by S.M.P.

## Acknowledgments

We thank Prof. Alessandro Quattrone, Alessia Soldano, Pierre Vanderhaeghen and members of CIBIO for helpful discussions; Giannino DelSal and Alessandra Rustighi for the pcDNA3-N1ICD-myc. The pMath1-creER plasmid was a gift from Robert Machold. We thank Sergio Robbiati (MOF facility) for supporting the work with mice models; Veronica De Sanctis and Roberto Bertorelli (NGS facility) for library generation and sequencing. This work was funded by a grant from The Giovanni Armenise-Harvard foundation (Career Development Award), My First AIRC Grant and CARITRO (to L.T.), program “Investissements d’avenir” ANR-10-IAIHU-06, the ICM foundation, Allen Distinguished Investigator Award and the Roger De Spoelberch Prize (to B.A.H.). F.A. is funded by Fondazione Umberto Veronesi post-doctoral fellowship and T.T.Z is supported by doctoral fellowships form the China Scholarship Council.

## Materials and methods

### Plasmids

The plasmid encoding a hyperactive form of the piggyBac transposase (pCMV HAhyPBase, pPBase) was a gift from the Wellcome Sanger Institute (Yusa K et al., 2011). The piggyBac donor plasmids pPB CAG MYC, pPB CAG Gfi1:FLAG-IRES-GFP, pPB CAG Otx2-IRES-GFP and pPB CAG Venus were previously described (Ballabio et al. 2020). These plasmids were used as backbones to insert before the start codon a loxP-STOP-loxP cassette by PCR, generating respectively pPB CAG LSL MYC, pPB CAG LSL Gfi1:FLAG-IRES-GFP and pPB CAG LSL Venus. The pPB CAG LSL N1ICD:V5-IRES-GFP plasmid was generated by substituting Gfi1:FLAG with the PCR-amplified N1ICD:V5 coding sequence derived from pcDNA3-N1ICD-myc (Rustighi et al., 2009). The pPB CAG LSL mCherry plasmid was generated by substituting Venus coding sequence with mCherry cDNA. The pMath1-creER plasmid was a gift from Robert Machold (Machold et al., 2005) and was used to generate the pMath1-cre plasmid, by substituting the creER coding sequence with the cre coding sequence. The pTbr2-cre plasmid was a gift from Tarik Haydar (Tyler et al., 2015). The pSox2-cre plasmid was generated by cloning the cre coding sequence into pGL3-Sox2 (Addgene Plasmid #101761). The pS100b-cre plasmid was generated by substituting the Sox2 promoter sequence from pSox2-cre with S100b promoter sequence, which was PCR-amplified from pEMS1384 (Addgene Plasmid #29304). The pHes5-d2EGFP plasmid was a gift from Ryoichiro Kageyama (Ohtsuka et al., 2006).

### Mice husbandry

Rosa26-LSL-MYC (#020458), Rosa26-LSL-tdTomato (#007908), Math1-creER (#007684), Ascl1-creER (#012882), Sox2-creER (#017593) were purchased from The Jackson Laboratory. CD1 mice were purchased from Charles River Laboratories. Mice were intraperitoneally injected with 75 mg/kg Tamoxifen at E9.5, E10.5, E13.5, P0, P1, P2, P5, P7 or P9. Animals were sacrificed at E15.5, P4, P7, P10, P14, P21, P30, P75 or at a humane endpoint as they displayed signs of morbidity. Mice were housed in a certified Animal Facility in accordance with European Guidelines. The experiments were approved by the Italian Ministry of Health as conforming to the relevant regulatory standards.

### *In vivo* transfection and *in utero* electroporation

For in vivo transfection, plasmid DNA and in vivo-jetPEI transfection reagent (Polyplus-transfection) were mixed according to manufacturer’s instructions. pPBase and piggyBac donor plasmids were mixed at a 1:4 ratio. The pPB CAG Venus or pPB CAG LSL Venus plasmids were always co-transfected as a reporter. P0-P2 mice were anesthetized on ice for 2 minutes, placed on a stage in a stereotactic apparatus and medially injected at lambda: −3.6 D/V: −1.6 with 4 µl of transfection mix using a pulled glass capillary and a FemtoJet microinjector (Eppendorf). *In utero* electroporation was performed at day E15.5, after defining day E0.5 by the observation of a vaginal plug. The pregnant female was anesthetized with 2% isoflurane and the uterine horns were exposed. After 1 µl solution (2.5 µg/µl) of plasmids in sterile saline was injected into the fourth ventricle of CD1 embryos, DNA was transferred using electric square pulses delivered by forceps-like electrodes (35 mV, pulse: 50 ms, pause: 950 ms, 5 pulses).

### Immunofluorescence and immunohistochemistry

Mice were intraventricularly perfused with 4% PFA, brains were dissected and post-fixed overnight in 4% PFA. Brains were either cryoprotected in 30% (w/v) sucrose in water and embedded in Frozen Section Compound (Leica, 3801480), or embedded in paraffin (brains were dehydrated with ethanol, then kept sequentially in xylene and paraffin solutions), or otherwise embedded in 5% agarose. Frozen Section Compound embedded brains were cryosectioned at 20-40 μm with a Leica CM 1850 UV Cryostat. Paraffin-embedded brains were sectioned using a Leica Microtome at 10 μm. Agarose-embedded tissues were sectioned using a Leica Vibratome at 100 μm. Immunofluorescence stainings were performed on glass slides. Blocking and Antibody solutions consisted of PBS supplemented with 3% goat serum, 0.3% Triton X-100 (Sigma). Primary antibodies were incubated overnight at 4°C and secondary antibodies for 1 hour at room temperature. Nuclei were stained with 1 µg/ml DAPI (Sigma). Sections and coverslips were mounted with Permanent Mounting Medium.

Immunohistochemistry stainings were performed on rehydrated paraffin sections. Antigen retrieval was performed by incubating slices for 30 min in retrieval solution (10 mM Sodium Citrate, 0.5% Tween-20 (v/v), pH 6.0) at 98°C. Primary antibodies were incubated overnight at 4°C and secondary antibodies for 1 hour at room temperature in Antibody solution. ABC solution was used 2 hours at room temperature (Vectastain Elite ABC Kit Standard PK-6100). The sections were incubated with the substrate at room temperature until suitable staining was observed (DAB Peroxidase Substrate Kit, SK-4100). Nuclei were counterstained with Hematoxylin.

The used antibodies are listed below:

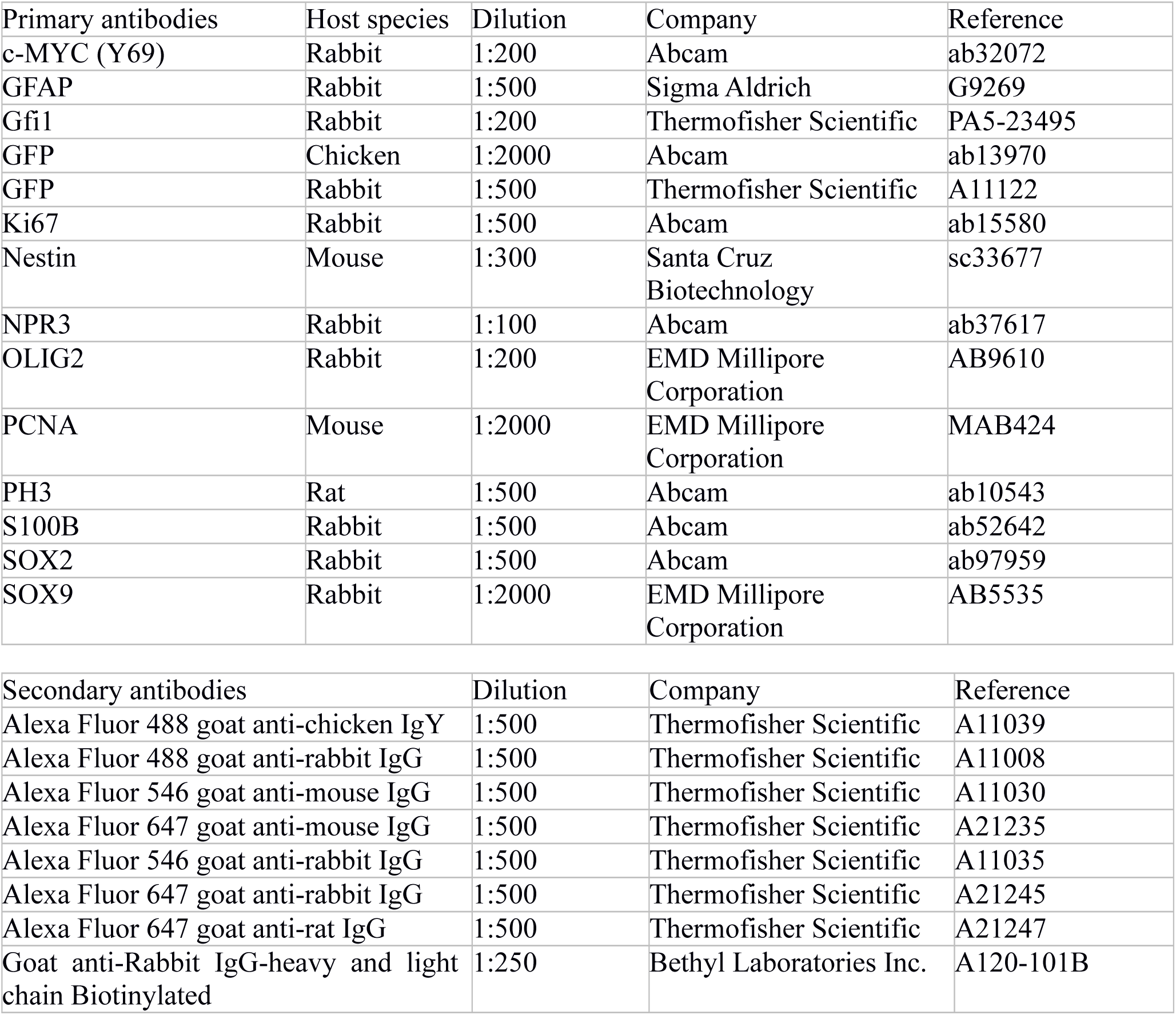

| Primary antibodies | Host species | Dilution | Company | Reference |
| --- | --- | --- | --- | --- |
| c-MYC (Y69) | Rabbit | 1:200 | Abcam | ab32072 |
| GFAP | Rabbit | 1:500 | Sigma Aldrich | G9269 |
| Gfi1 | Rabbit | 1:200 | Thermofisher Scientific | PA5-23495 |
| GFP | Chicken | 1:2000 | Abcam | ab13970 |
| GFP | Rabbit | 1:500 | Thermofisher Scientific | A11122 |
| Ki67 | Rabbit | 1:500 | Abcam | ab15580 |
| Nestin | Mouse | 1:300 | Santa Cruz<br>Biotechnology | sc33677 |
| NPR3 | Rabbit | 1:100 | Abcam | ab37617 |
| OLIG2 | Rabbit | 1:200 | EMD Millipore<br>Corporation | AB9610 |
| PCNA | Mouse | 1:2000 | EMD Millipore<br>Corporation | MAB424 |
| PH3 | Rat | 1:500 | Abcam | ab10543 |
| S100B | Rabbit | 1:500 | Abcam | ab52642 |
| SOX2 | Rabbit | 1:500 | Abcam | ab97959 |
| SOX9 | Rabbit | 1:2000 | EMD Millipore<br>Corporation | AB5535 |

| Secondary antibodies | Dilution | Company | Reference |
| --- | --- | --- | --- |
| Alexa Fluor 488 goat anti-chicken IgY | 1:500 | Thermofisher Scientific | A11039 |
| Alexa Fluor 488 goat anti-rabbit IgG | 1:500 | Thermofisher Scientific | A11008 |
| Alexa Fluor 546 goat anti-mouse IgG | 1:500 | Thermofisher Scientific | A11030 |
| Alexa Fluor 647 goat anti-mouse IgG | 1:500 | Thermofisher Scientific | A21235 |
| Alexa Fluor 546 goat anti-rabbit IgG | 1:500 | Thermofisher Scientific | A11035 |
| Alexa Fluor 647 goat anti-rabbit IgG | 1:500 | Thermofisher Scientific | A21245 |
| Alexa Fluor 647 goat anti-rat IgG | 1:500 | Thermofisher Scientific | A21247 |
| Goat anti-Rabbit IgG-heavy and light chain Biotinylated | 1:250 | Bethyl Laboratories Inc. | A120-101B |

### Imaging

Images were acquired with a Zeiss Axio Imager M2 (Axiocam MRc, Axiocam MRm); confocal imaging was performed by Leica TCS Sp5 or Leica TCS Sp8; live imaging was performed by Nikon TI2 equipped with Spinning Disc X-light V2. Images were processed using ImageJ software.

### Organoids maintenance and modification

Human iPS cells (iPSCs, ATCC-DYS0100) were maintained in self renewal on a layer of Geltrex (Gibco, A14133-01), in Essential 8 Basal Medium (Gibco, A15169-01) supplemented with E8 Supplement (50X, Gibco A15171-01) and P/S (Pen 100 Units/ml, Strep 100 μg/ml, GIBCO, 15140-122). All cells were mycoplasma free. iPSCs were dissociated with EDTA (Invitrogen, 15575-038) 0.5mM, pH 8.0, for 3 minutes incubation, to maintain cell clusters. Cerebellar organoids were cultured as described in Muguruma et al., 2015; Ishida et al., 2016 and were electroporated at 35 days of differentiation protocol with pCMV HAhyPBase (pPBase), pPB CAG LSL Venus, pPB CAG LSL Gfi1:FLAG-IRES-GFP, pPB CAG LSL MYC and either pS100b-cre (S100b-cre, LSL-GM), or pSox2-cre (Sox2-cre, LSL-GM) resuspended in Buffer 5 (under patent, Ballabio et al., 2020). For the 24 hours analysis, organoids were electroporated at 35 days of differentiation protocol with pPB CAG LSL Venus and either pS100b-cre (S100b-cre, LSL-V) or pSox2-cre (Sox2-cre, LSL-V). Organoids were transferred inside the electroporation cuvettes (VWR, ECN 732-1136, 2mm) and electroporation was performed with the Gene Pulser XcellTM. 24 hours after electroporation for S100b-cre, LSL-V and Sox2-cre, LSL-V organoids and 39 days after electroporation for S100b-cre, LSL-GM, they were fixed in 4% PFA, cryoprotected in 20% sucrose and embedded in Frozen Section Compound (Leica, 3801480). Organoids were cryosectioned at 40 μm with Thermo Scientific HM525 NX cryostat.

### Cell quantification and statistical analysis

Cell quantification in P4 CD1 mouse cerebella transfected with pS100b-cre + pPB-LSL-Venus: Venus^+^ cells coming from at least 2 mice were analyzed for each quantification. All data are presented as mean + S.E.M.

Cell quantification in P4 CD1 mouse cerebella transfected with pS100b-cre + pPB-LSL-mCherry + pHes5-d2EGFP or pMath1-cre + pPB-LSL-mCherry + pHes5-d2EGFP: Venus^+^ cells coming from at least 3 mice were analyzed for each quantification. All data are presented as mean + S.E.M. Data were compared using an unpaired Student’s t test, two-tailed.

Organoids quantitative 24 hours analysis: data are presented as mean + s.e.m., for each condition 10-17 organoids were examined and at least 30-45 cells were quantified.

Live clustering analysis: brightfield and fluorescence images of the electroporated organoids were acquired and Venus^+^ fluorescent area was quantified using ImageJ. Groups of Venus ^+^ cells with an area ≥ 3500 μm^2^ were considered clusters.

### RNA extraction, library generation and sequencing

Total RNA was extracted from fresh-frozen tumor tissues with TRIzol Reagent (Invitrogen). Libraries were prepared from the extracted RNAs using the QuantSeq 3’mRNA-Seq Library Prep Kit-FWD (Lexogen, Vienna, Austria) using 0.5-1 μg of RNA per library and following the manufacturers’ instructions. We modified the standard protocol adding Unique Molecular Identifiers (UMI) during the second strand synthesis step. Indices from the Lexogen i7 6nt Index Set and i5 6nt Dual Indexing Add-on Kits (Cat. No. 044.96 and 047.96, Lexogen) were used, and 15-19 cycles of library amplification were performed. Libraries were eluted in 30 μL of the kit’s Elution Buffer. The double-stranded DNA concentration was quantified using the Qubit dsDNA HS Assay Kit (ThermoFisher), ranging from 3 to 12 ng/μL. The molar concentration of cDNA molecules in the individual libraries was calculated from the double-stranded DNA concentration and the single library average size (determined on a PerkinElmer Labchip GX). An equal number of cDNA molecules from each library were pooled and the final pool was purified once more in order to remove any free primer and prevent index-hopping. The pooled libraries were sequenced in a Novaseq 6000 instrument (Illumina, San Diego, CA) on an SP flowcell, producing 900M single reads 100nt and in a Hiseq2500 (Illumina, San Diego, CA) on a rapid run lane, producing 166M single reads 100nt.

### Mouse cerebellum single cell RNA-sequencing data integration

The full cell type annotation and the gene expression counts per sample were obtained from the shared materials of the corresponding mice cerebellum single-cell study (*Carter et al. 2018*). The samples on time-points P0 and P4 were combined and re-analyzed using Seurat v2.3.2 package (*Butler et al. 2018*). Further, the cells from 3 selected assigned clusters α,β and γ were split into groups based on the presence of either Math1 or S100b expression. Afterwards, the differential gene expression comparison among these groups (Math1^+^ vs S100b^+^ cluster α Math1^+^ vs S100b^+^ cluster β, Math1^+^ vs S100b^+^ cluster γ) was performed using t-test separately on time points P0 and P4. Finally, adjusted p-values were computed using qvalue R package.

### Mouse model gene expression analysis

The RNA-sequencing transcripome profiling was performed on the mouse model materials in the corresponding cohorts: pPB-CAG-MYC + pPB-CAG-Gfi1 (n=2), pPB-CAG-MYC + pPB-CAG-Otx2 (n=3) and pMath1-cre + pPB-LSL-Myc + pPB-LSL-Gfi1 + pPB-LSL-N1ICD (n=3). The reads were aligned using STAR v2.4.1 tool *(Dobin et al., 2013*) to mm10 reference genome, and gene expression counts were computed using featureCounts module of the Subread package v1.4.6 *(Liao et al., 2014*) with Ensembl GRCm38 v72 annotation. The quality control was performed with Qualimap v2 using RNA-seq QC mode (*Okonechnikov et al., 2016*). Further, for comparison to human tumors, RNA-sequencing gene expression data was collected from medulloblastoma landscape study (*Northcott et al 2017)*. The orthologus genes common between species were filtered using biomart R package and corresponding Ensembl database annotation. Unsupervised clustering was performed on combined RNA-seq cohorts after Combat batch effect adjustment (*Leek et al, 2012*) with focus on 500 top most highly variable genes.

---

**Supplementary Figure 1.**
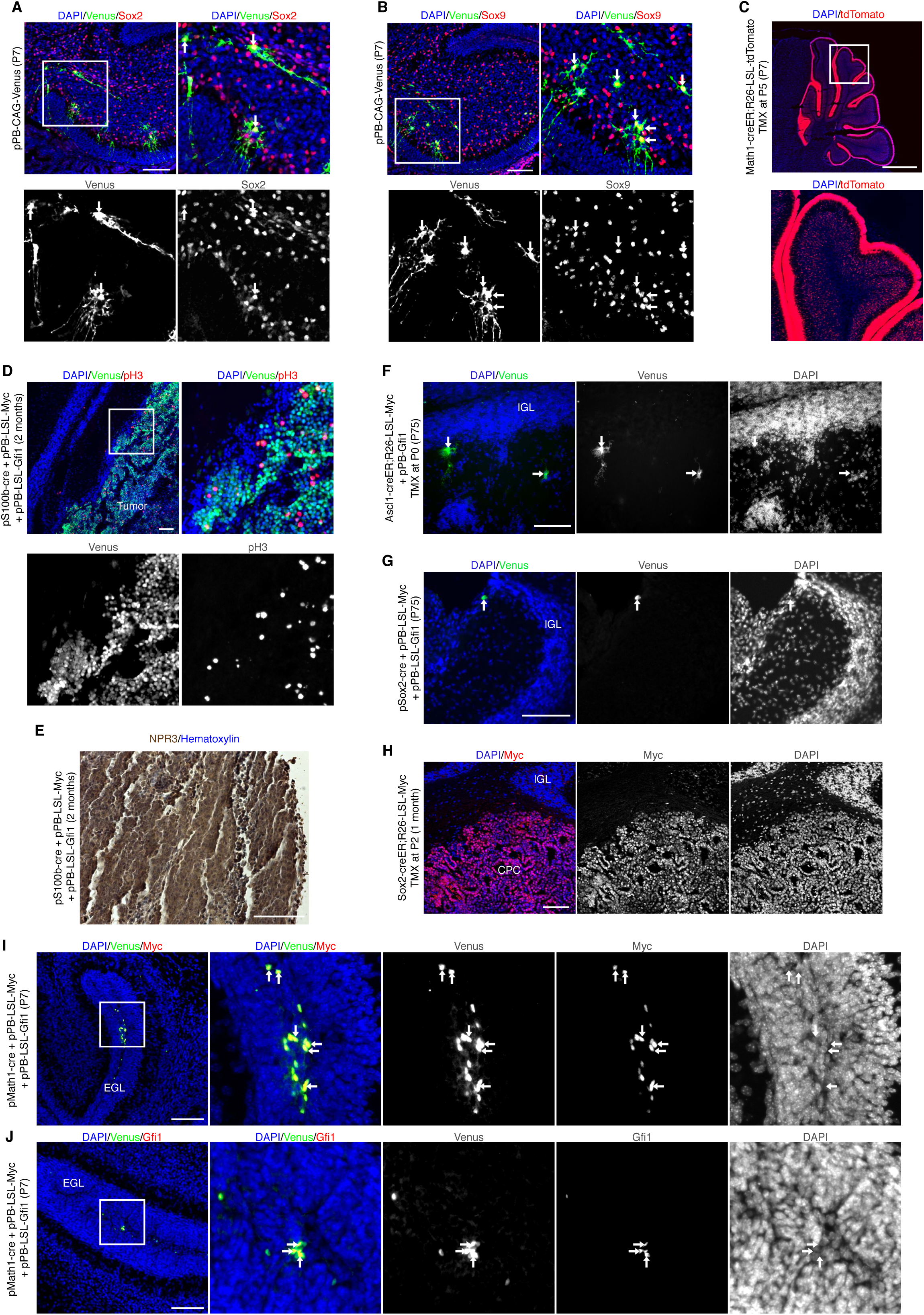
Efficient and stable expression of transgenes in mouse cerebellum. (**A,B**) Confocal images of Venus and Sox2 (**A**), Venus and Sox9 (**B**) immunofluorescence of CD1 cerebellum 7 days after transfection with pPBase and pPB-CAG-Venus at P0. The white squares in (**A,B**) mark the regions shown at higher magnification. Arrows point to double positive cells. (**C**) DAPI staining and tdTomato fluorescence of P7 Math1-creER;R26-LSL-tdTomato mouse cerebellum injected with tamoxifen at P5. The white square marks the region shown at higher magnification. (**D**) Confocal images of Venus and pH3 immunofluorescence of tumors in CD1 mice 2 months after transfection with pPBase + pS100b-cre + pPB-LSL-Myc + pPB-LSL-Gfi1 + pPB-LSL-Venus at P0. The white square marks the region shown at higher magnification. (**E**) Immunohistochemistry for NPR3 and hematoxylin staining in tumors of CD1 mice after transfection with pPBase + pS100b-cre + pPB-LSL-Myc + pPB-LSL-Gfi1 + pPB-LSL-Venus at P0. (**F**) DAPI staining and Venus immunofluorescence of sagittal brain sections of Ascl1-creER;R26-LSL-Myc mouse 2.5 months after transfection with pPBase + pPB-Gfi1 + pPB-Venus at P0 and tamoxifen injection at P0. Arrows point to Venus^+^ cells. (**G**) DAPI staining and Venus immunofluorescence of sagittal brain sections of CD1 mouse cerebellum 2.5 months after transfection with pPBase + pSox2-cre + pPB-LSL-Myc + pPB-LSL-Gfi1 + pPB-LSL-Venus at P0. Arrows point to Venus^+^ cells. (**H**) Confocal images of Myc immunofluorescence in choroid plexus carcinoma (CPC) of Sox2-creER;R26-LSL-Myc mouse 1 month after tamoxifen injection at P2. (**I,J**) Confocal images of Venus and Myc (**I**), Venus and Gfi1 (**J**) immunofluorescence in CD1 mouse cerebella 7 days after transfection with pPBase + pMath1-cre + pPB-LSL-Myc + pPB-LSL-Gfi1 + pPB-LSL-Venus at P0. The white squares in (**I,J**) mark the regions shown at higher magnification. Arrows point to double-positive cells. Scale bars: 100 µm in (**A,B,D-J**), 1 mm in (**C**).

**Supplementary Figure 2.**
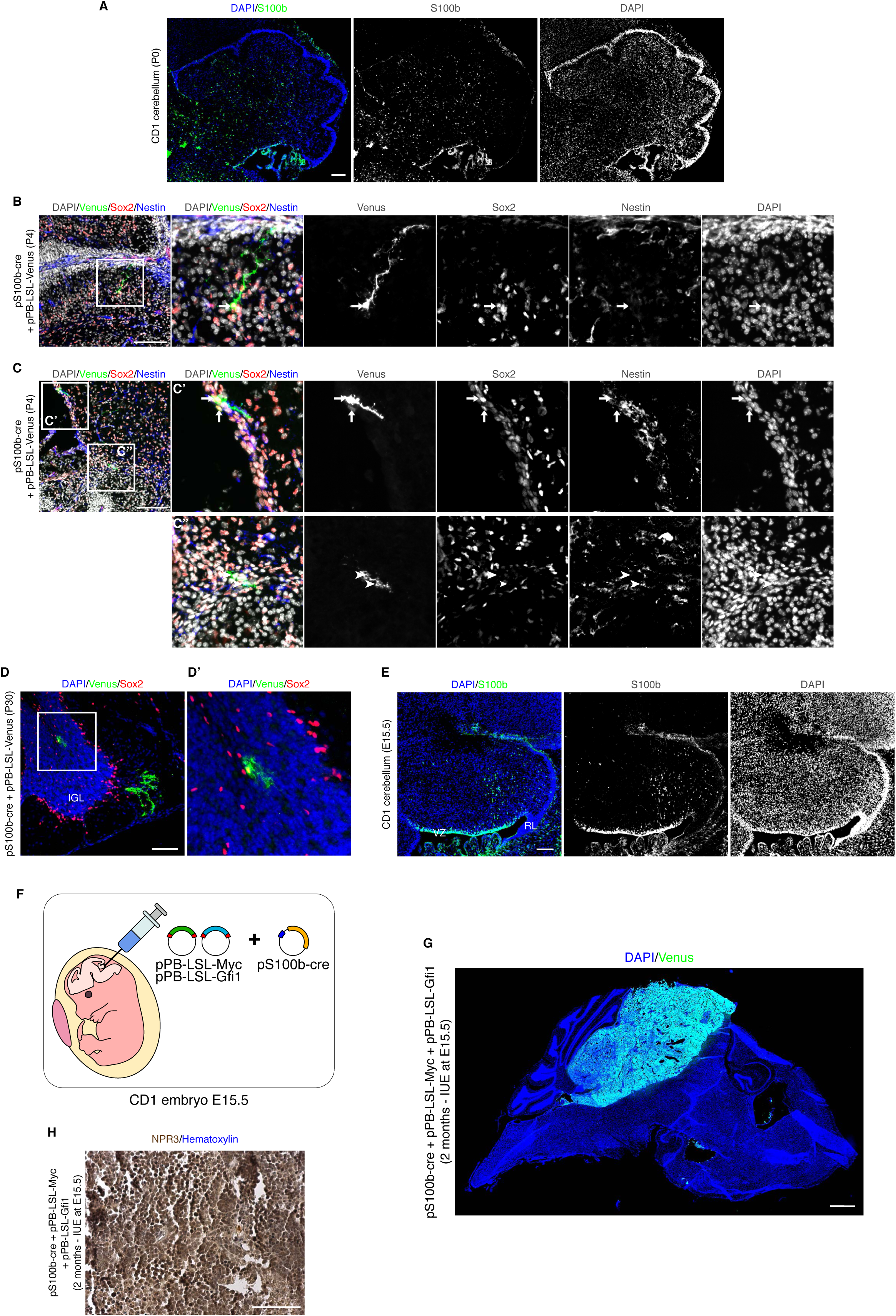
S100b^+^ cells during postnatal and embryonic cerebellar development. (**A**) Confocal images of S100b immunofluorescence of CD1 mouse cerebellum at P0. (**B,C**) Confocal images of Venus, Sox2 and Nestin immunofluorescence of CD1 mouse cerebellum 4 days after transfection with pPBase + pS100b-cre + pPB-LSL-Venus at P0. The white squares mark the regions shown at higher magnification. Arrows point to Venus^+^/Sox2^+^ cells, whereas arrowheads point to Venus^+^/Sox2^-^cells. (**D**) Confocal images of Venus and Sox2 immunofluorescence of CD1 mouse cerebellum 30 days after transfection with pPBase + pS100b-cre + pPB-LSL-Venus at P0. The white square in (**D**) marks the region shown at higher magnification in (**D’**). (**E**) Confocal images of S100b immunofluorescence of CD1 mouse cerebellum at E15.5. VZ: ventricular zone; RL: rhombic lip. (**F**) Schematic representation of *in utero* electroporation to overexpress Myc and Gfi1 in S100b^+^ cells at E15.5. (**G**) DAPI staining and Venus immunofluorescence of sagittal brain section of CD1 mouse (2 months) after *in utero* electroporation with pPBase + pS100b-cre + pPB-LSL-Myc + pPB-LSL-Gfi1 + pPB-LSL-Venus at E15.5. (**H**) Immunohistochemistry for NPR3 and hematoxylin staining in tumors of CD1 mice after *in utero* electroporation with pPBase + pS100b-cre + pPB-LSL-Myc + pPB-LSL-Gfi1 + pPB-LSL-Venus at E15.5. Scale bars: 1 mm in (**G**), 100 µm in (**A-E,H**).

**Supplementary Figure 3.**
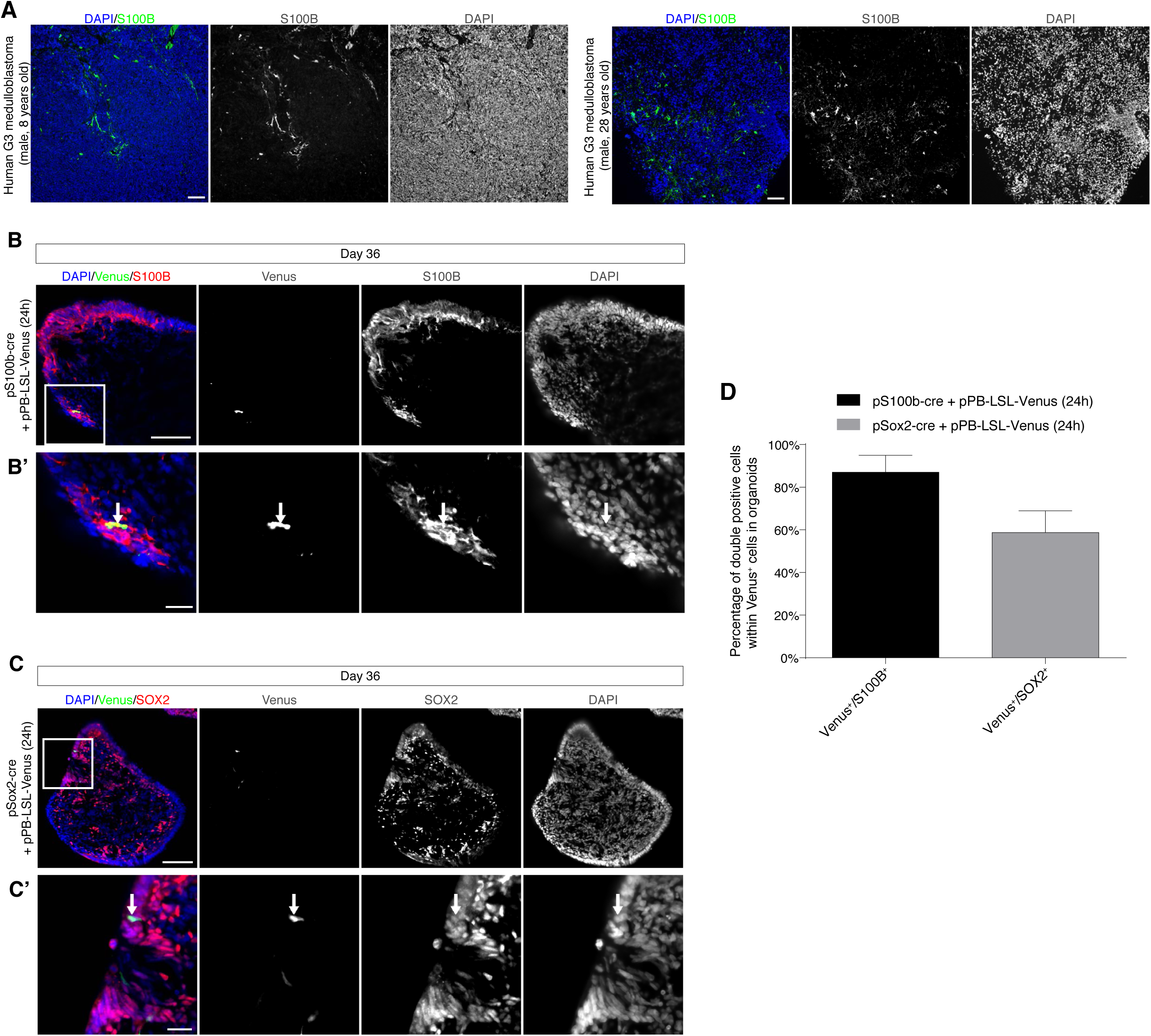
Lineage tracing in human cerebellar organoids electroporated with pS100b-cre and pSox2-cre. **(A)** Confocal images of S100B immunofluorescence of human Group 3 medulloblastoma samples from an 8-year-old male patient and a 28-year-old male patient. **(B)** Confocal images of Venus and S100B immunofluorescence of cerebellar organoids at day 36, electroporated at day 35 with pS100b-cre + pPB-LSL-Venus. The white square in (**B**) marks the region shown at higher magnification in (**B’**). (**C**) Confocal images of Venus and Sox2 immunofluorescence of cerebellar organoids at day 36, electroporated at day 35 with pSox2-cre + pPB-LSL-Venus. The white square in (**C**) marks the region shown at higher magnification in (**C’**). Arrows indicate double positive cells. (**D**) Quantification of cerebellar organoids at day 36, electroporated at day 35 with either pS100b-cre + pPB-LSL-Venus (n=17 biologically independent organoids) or pSox2-cre + pPB-LSL-Venus (n=10 biologically independent organoids). Arrows indicate double positive cells. Scale bars: 100 µm in (**A-C**), 25 µm in (**B’,C’**).

**Supplementary Figure 4.**
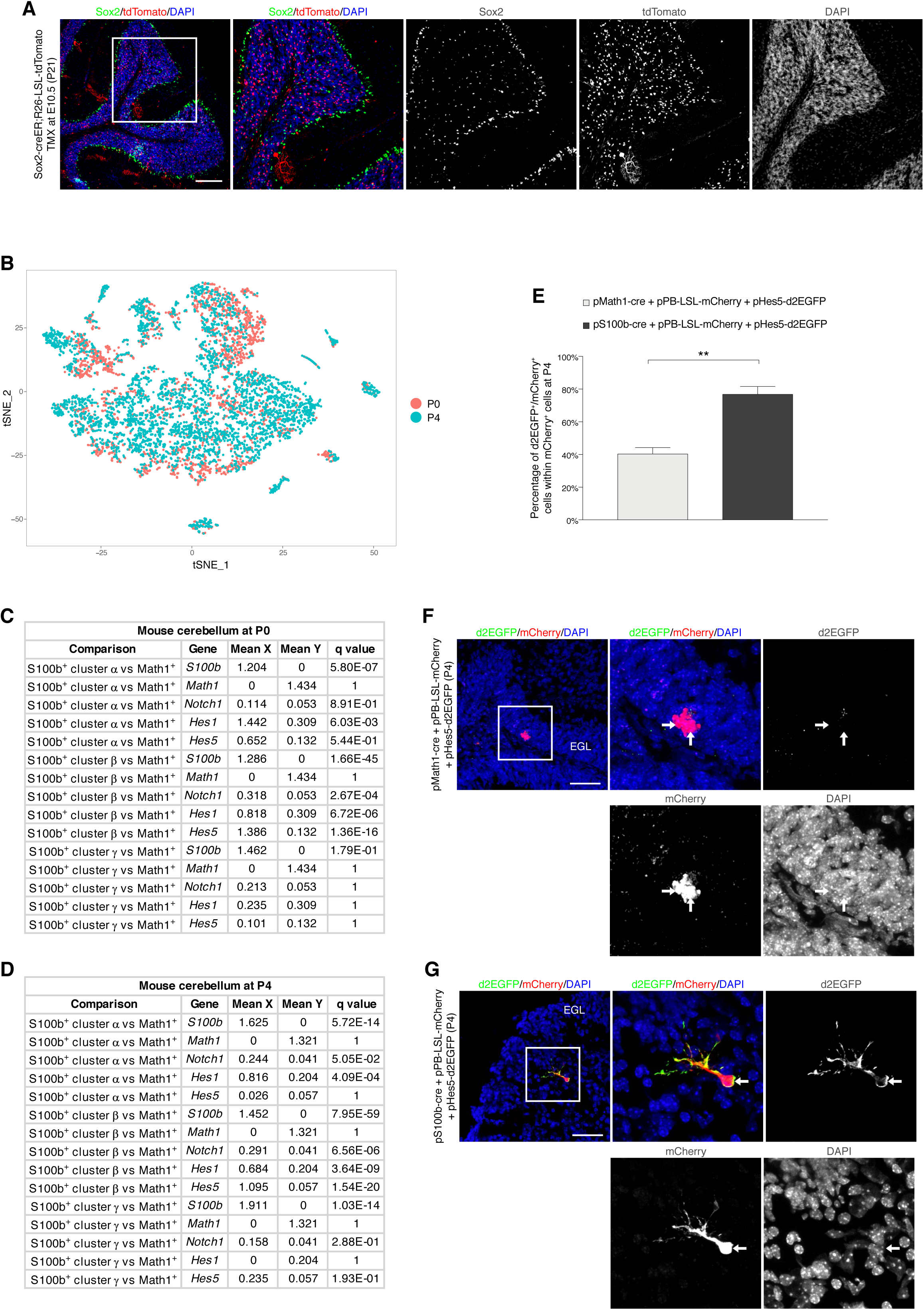
Notch pathway activation levels in different cell lineages. (**A**) Confocal images of Sox2 and tdTomato immunofluorescence on Sox2-creER;R26-LSL-tdTomato mouse cerebellum at P21, after tamoxifen injection at E10.5. (**B**) t-distributed stochastic neighbor embedding (t-SNE) visualizations of cerebellar derived developmental time points at P0 and P4. Each point represents one cell, colors represent the time points. (**C,D**) Differential expression analysis results in selected scRNAseq subclusters between S100b, Math1, Notch1, Hes, Hes5 at P0 (**C**) and P4 (**D**) in four groups: Math1^+^ cells, S100b^+^ cells within cluster α, S100b^+^ cells within cluster β and S100b^+^ cells within cluster γ. (**E**) Quantification of d2EGFP^+^/mCherry^+^ within mCherry^+^ cells in CD1 mouse cerebellum 4 days after transfection with either pPBase + pMath1-cre + pPB-LSL-mCherry + pHes5-d2EGFP (n=4 mice, 77 cells) or pPBase + pS100b-cre + pPB-LSL-mCherry + pHes5-d2EGFP (n=3 mice, 99 cells) at P0. (**F**) Confocal images of mCherry and d2EGFP immunofluorescence of transfected cells in CD1 mouse 4 days after transfection with pPBase + pMath1-cre + pPB-LSL-mCherry + pHes5-d2EGFP at P0. Arrows point to mCherry^+^/d2EGFP^-^cells. (**G**) Confocal images of mCherry and d2EGFP immunofluorescence of transfected cells in CD1 mouse 4 days after transfection with pPBase + pS100b-cre + pPB-LSL-mCherry + pHes5-d2EGFP at P0. Arrow points to a mCherry^+^/d2EGFP^+^ cell. The white squares in the top-left panels (**A,F,G**) mark the regions shown at higher magnification. Error bars in (**E**) represent s.e.m. Unpaired Student’s t test, two-tailed: **p-value < 0.01. Scale bars: 200 µm in (**A**), 50 µm in (**F,G**).

**Supplementary Figure 5.**
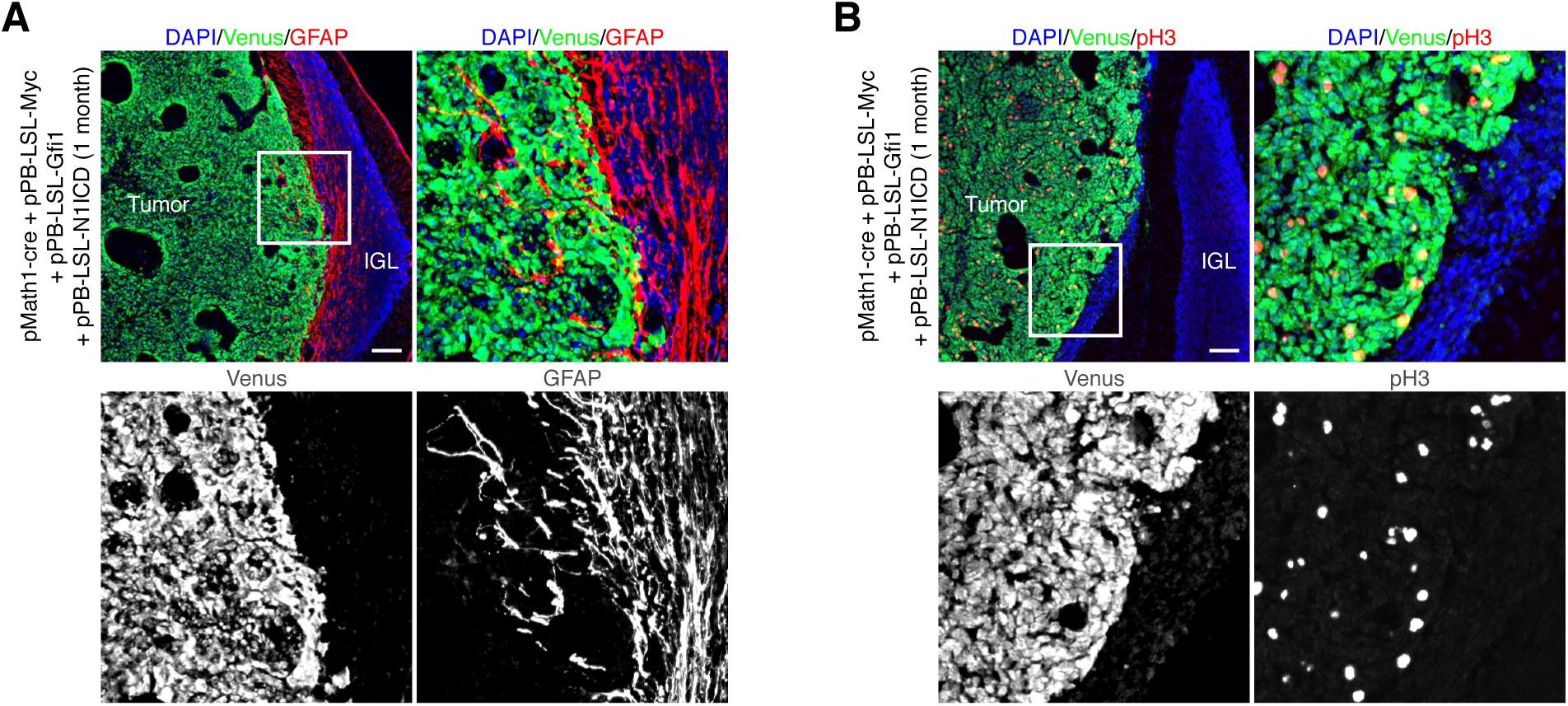
Expression of N1ICD, MYC and Gfi1 in Math1^+^ cells induces Group 3 MB. (**A,B**) Confocal images of GFP (Venus) and GFAP (**A**) or pH3 (**B**) immunofluorescence of tumors in CD1 mouse 1 month after transfection with pPBase ^+^ pMath1-cre + pPB-LSL-Myc + pPB-LSL-Gfi1 + pPB-LSL-N1ICD + pPB-LSL-Venus at P0. The white squares in the top-left panels mark the regions shown at higher magnification. Scale bars: 100 µm in (**A,B**).

**Table S1.** Tumor incidence upon *in vivo* transfection of cerebellar cells. Number of mice developing tumors within mice showing transfected cerebellar cells upon injection at P0. TMX: Tamoxifen administration. CPC: Choroid Plexus Carcinoma.

| Tumor incidence upon injection in transgenic mice |  |  |  |  |
| --- | --- | --- | --- | --- |
|  | TMX | R26-LSL-MYC | R26-LSL-MYC + pPB-CAG-Gfi1 | + pPB-LSL-MYC + pPB-LSL-Gfi1 |
| Math1-creER | P2 |  | 0/10 | 0/8 |
|  | P9 |  | 0/6 |  |
| Ascl1-creER | P0 |  | 0/3 |  |
|  | P2 |  | 0/9 | 0/6 |
|  | P7 |  | 0/6 |  |
| Sox2-creER | P2 | 6/6 (CPC) | 1/1 (CPC) |  |

| Tumor incidence upon injection in CD1 mice |  |  |
| --- | --- | --- |
|  | TMX | + pPB-LSL-MYC + pPB-LSL-Gfi1 |
| pMath1-creER | P1 | 0/10 |
| pMath1-cre |  | 0/10 |
| pTbr2-cre |  | 0/10 |
| pS100b-cre |  | 3/16 |
| pSox2-cre |  | 0/8 |

